# Regional interneuron transcriptional changes reveal pathologic markers of disease progression in a mouse model of Alzheimer’s disease

**DOI:** 10.1101/2023.11.01.565165

**Authors:** Kevin S. Chen, Mohamed H. Noureldein, Diana M. Rigan, John M. Hayes, Masha G. Savelieff, Eva L. Feldman

**Author notes:** Correspondence: Eva L. Feldman, MD, PhD, 5017 AAT-BSRB, 109 Zina Pitcher Place, Ann Arbor, MI 48109. Equally contributing authors (Kevin S. Chen, Mohamed H. Noureldein).

## Abstract

Alzheimer’s disease (AD) is a progressive neurodegenerative disorder and leading cause of dementia, characterized by neuronal and synapse loss, amyloid-β and tau protein aggregates, and a multifactorial pathology involving neuroinflammation, vascular dysfunction, and disrupted metabolism. Additionally, there is growing evidence of imbalance between neuronal excitation and inhibition in the AD brain secondary to dysfunction of parvalbumin (PV)- and somatostatin (SST)-positive interneurons, which differentially modulate neuronal activity. Importantly, impaired interneuron activity in AD may occur upstream of amyloid-β pathology rendering it a potential therapeutic target. To determine the underlying pathologic processes involved in interneuron dysfunction, we spatially profiled the brain transcriptome of the 5XFAD AD mouse model versus controls, across four brain regions, dentate gyrus, hippocampal CA1 and CA3, and cortex, at early-stage (12 weeks-of-age) and late-stage (30 weeks-of-age) disease. Global comparison of differentially expressed genes (DEGs) followed by enrichment analysis of 5XFAD versus control highlighted various biological pathways related to RNA and protein processing, transport, and clearance in early-stage disease and neurodegeneration pathways at late-stage disease. Early-stage DEGs examination found shared, *e.g*., RNA and protein biology, and distinct, *e.g*., N-glycan biosynthesis, pathways enriched in PV-versus somatostatin SST-positive interneurons and in excitatory neurons, which expressed neurodegenerative and axon- and synapse-related pathways. At late-stage disease, PV-positive interneurons featured cancer and cancer signaling pathways along with neuronal and synapse pathways, whereas SST-positive interneurons showcased glycan biosynthesis and various infection pathways. Late-state excitatory neurons were primarily characterized by neurodegenerative pathways. These fine-grained transcriptomic profiles for PV- and SST-positive interneurons in a time- and spatial-dependent manner offer new insight into potential AD pathophysiology and therapeutic targets.

## INTRODUCTION

Alzheimer’s disease (AD) is a progressive neurodegenerative disorder causing memory loss, learning deficits, and a gradual inability to perform activities of daily living.^1^ The illness is characterized by neuronal and synapse loss, especially in the hippocampus and entorhinal cortex, which eventually spreads to the entire cerebral cortex.^2^ In tandem with brain atrophy, plaques of amyloid-beta (Aβ) and neurofibrillary tangles of tau accumulate^3^ along with a multifactorial pathology involving neuroinflammation,^4, 5^ vascular dysfunction, and disrupted metabolism.^6^

In addition to neuronal and synapse loss, AD patients demonstrate an imbalance between neuronal excitation and inhibition in the brain, leading to hyperexcitability mediated through a breakdown in inhibitory interneuron activity.^7–9^ Interneuron dysfunction in AD suppresses neurogenesis and GABAergic transmission^10^ impairs learning and cognition^11^ and precedes major clinical manifestations of AD.^12–17^ Therefore, interneuron pathology in AD could feasibly occur upstream of Aβ pathology. Despite decades of research, AD remains incurable, and patients inevitably suffer progressive cognitive decline and death. The vast majority of therapeutics have targeted Aβ^18^ and have been largely unsuccessful, necessitating novel approaches. As an early pathology possibly occurring upstream of Aβ pathology, GABA interneurons may serve as a gateway to novel AD therapeutics and therapeutic targets. Indeed, interneuron transplant^19^ or stimulated activity^20^ in AD animal models restores brain rhythms and cognition. Still, the underlying pathologic processes involved in interneuron dysfunction in AD remain unknown.

Interneurons normally regulate excitatory neuronal signaling and tune neural networks during the performance of memory- and cognition-dependent tasks.^21^ Moreover, proper interneuron activity clears Aβ,^20, 22^ whereas neural network hyperexcitability elevates Aβ levels.^23, 24^ In a vicious cycle, raised Aβ levels are toxic to and interfere with interneuron activity;^9, 11^ thus, as anticipated, the AD brain is characterized by abnormal synaptic transmission, aberrant neural network activity, and a loss of interneurons in both humans and mouse models.^9, 11^ Interneurons are classified into five subtypes based on differential molecular expression; specifically in AD pathophysiology, interneurons expressing parvalbumin (PV) and somatostatin (SST) are the main modulators of cortical and hippocampal circuits.^9, 21^ Systematic analysis of the literature suggests PV interneurons may be impacted earlier in the disease course whereas there is variability in the timeframe affecting SST interneurons.^21^ Nevertheless, as early pathological events, interneurons are of vital interest as possible cellular therapeutics^19^ and/or targets.^20^

Advancing possible interneuron applications requires fine-grained molecular information *in situ* in the brain. However, despite the pioneering work on interneuron dysfunction in AD, the transcriptomic changes occurring temporally and spatially in interneurons during disease progression have not been comprehensively described.^21^ This information could help identify an optimal timeframe and potential niches for interneuron transplant. Moreover, interneurons likely behave differently at various disease stages and possibly across anatomic subregions of the brain; however, few studies have described the exact mechanisms initiating interneuron dysfunction. Herein, using a spatial platform, we cataloged gene expression changes in interneurons and excitatory neurons in the brains of 5XFAD AD mice versus non-carrier controls at early-stage (12 weeks-of-age) and late-stage (30 weeks-of-age) disease. We analyzed differentially expressed genes (DEGs) followed by pathway enrichment analysis in 5XFAD versus control at the global level (early- versus late-stage), neuron-specific level (Pv-, Sst-, and excitatory-specific DEGs), and in a region-dependent manner (dentate gyrus [DG], hippocampal CA1 and CA3, and cortex). Overall, for the first time to our knowledge, we provide fine-grained transcriptomic profiles for Pv- and Sst-positive interneurons in a time- and spatial-dependent manner.

## METHODS AND MATERIALS

### Animal model

Animal procedures were approved by the University of Michigan Institutional Animal Care and Use Committee (Protocol #PRO00010247). 5XFAD male mice (catalog # 034848) and non-carrier male littermates were obtained from The Jackson Laboratory (Bar Harbor, ME). 5XFAD mice express five familial AD mutations in total, three to human amyloid-beta precursor protein [APP; Swedish (K670N/M671L), Florida (I716V), London (V717I)] and two to human presenilin-1 (PS1; M146L, L286V). Mice were maintained in cages with littermates in a specific-pathogen-free facility following a 12/12-h light/dark at 20 ± 2 °C. The facility was maintained by the Unit for Laboratory Animal Medicine at the University of Michigan, which also monitored the health of the animals daily. At 12 wk (early disease stage, presymptomatic; n=3 5XFAD, n=3 controls) and 30 wk of age (late disease stage, symptomatic; n=3 5XFAD, n=3 controls), mice were sacrificed by intraperitoneal pentobarbital injection (Fatal-Plus, Vortech Pharmaceuticals, Dearborn, MI) followed by perfusion with phosphate buffered saline. Brains were removed from the calvarium and dissected along the midline sagittal plane. One brain hemisphere was fixed in 4% paraformaldehyde (PFA) for 24 h, cryoprotected in escalating sucrose gradients, embedded, and cryosectioned in coronal plane sections (10 µm thickness) onto histologic slides and stored at –80 °C. Every tenth slide was stained by hematoxylin/eosin to identify brain structures.

### In situ hybridization to identify neuron subtypes for spatial transcriptomics

FISH was performed on brain sections using RNAscope probes (Advanced Cell Diagnostics, Bio-Techne, Newark, CA) to identify interneuron subtypes. After post-fixation in 4% PFA, dehydration/antigen retrieval/protease treatment per manufacturer protocol (with omission of hydrogen peroxide), Pv-positive interneurons were stained using a Pvalb probe (#421931-C2) developed using Opal dye 620 (Akoya Biosciences, Marlborough, MA, # FP1495001KT), Sst-positive interneurons were stained using a Sst probe (#404631-C3) with Opal dye 570 (#FP1488001KT), and Vglut-positive excitatory neurons were stained using a VGluT probe (#481851) with Opal dye 690 (# FP1497001KT). Nuclei were stained with SYTO 13 (Thermo Fisher Scientific, Waltham, MA) and slides stored in phosphate buffered saline and submitted the same day for spatial transcriptomics.

### Spatial transcriptomics and selecting regions of interest (ROIs)

Spatial transcriptomics was performed by GeoMx DSP (NanoString, Seattle, WA), a platform for spatially analyzing gene expression in tissue in a highly multiplexed manner.^25^ Four slides in total were submitted for spatial transcriptomics. Two slides were for early-stage analysis and were duplicates of 4 brain sections (n=2 5XFAD 12 wk, n=2 controls 12 wk); two slides were for late-stage analysis and were duplicates of 4 brain sections (n=2 5XFAD 30 wk, n=2 controls 30 wk). Duplicates were submitted to account for potential inter-slide variation. Briefly, slides are hybridized to a master mix of barcoded oligonucleotide probes directed to the whole mouse transcriptome. Regions of interest (ROIs) were defined in the DG, CA1, CA3, and cortex of the brain (**Figure 1**). Image segmentation permitted identification of interneurons defined by Pv- and Sst-positive staining, whereas excitatory neurons were identified by Vglut-positive staining.

**Figure 1.**
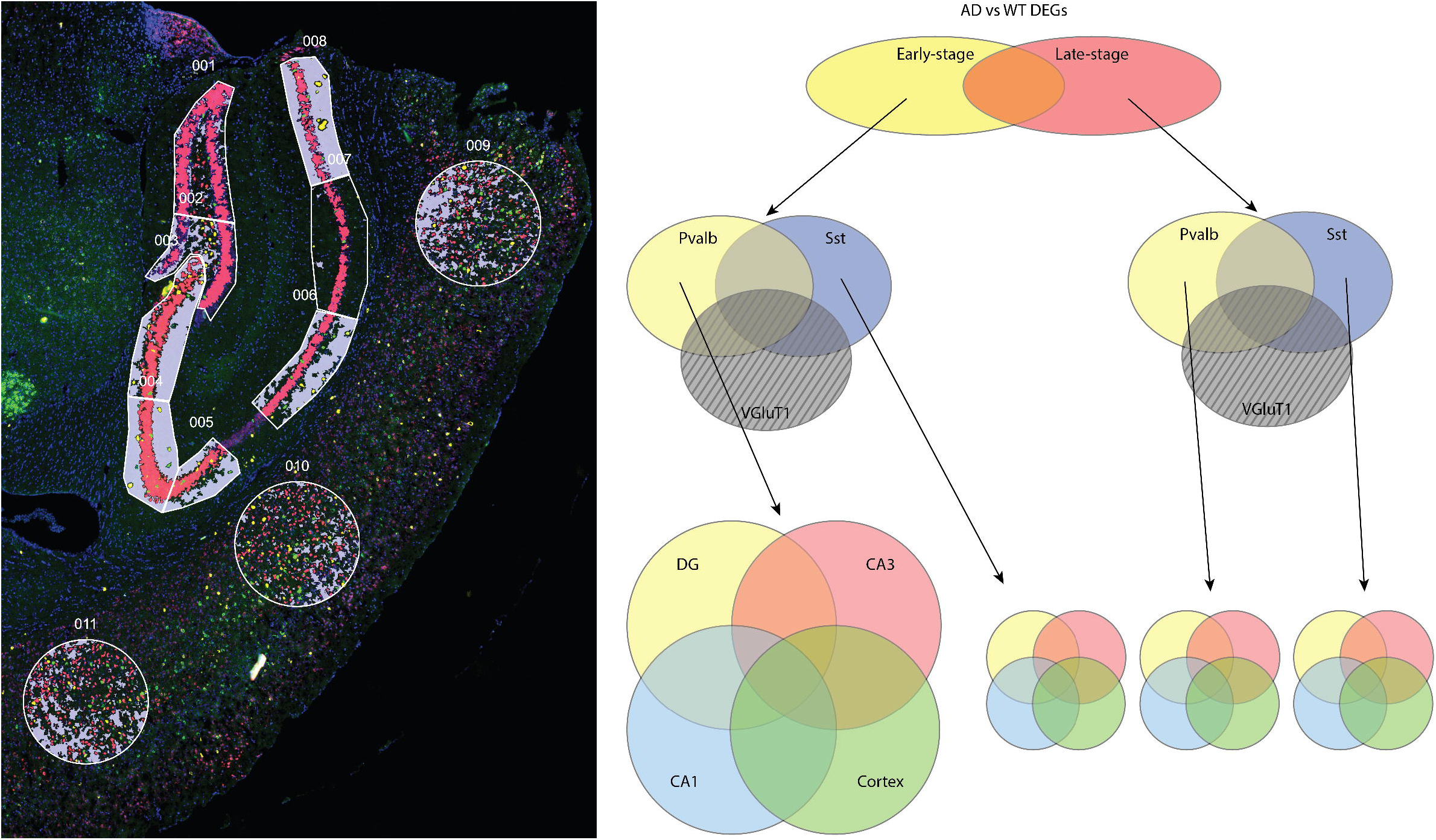
Spatial transcriptomics of mouse brain and study scheme. (**A**) Representative image of a mouse brain section stained by FISH and analyzed by spatial transcriptomics. Image segmentation within regions of interest (ROIs) during spatial transcriptomics captures whole mouse transcriptome from specific cell types: Pv- (yellow) and Sst- (green) positive interneurons amidst Vglut-positive excitatory neurons (red) against a background of non-neuronal cells (purple). ROIs capture transcriptional levels within defined anatomic subregions: dentate gyrus (DG; 001, 002), hippocampal CA3 (003, 004, 005) and CA1 (006, 007, 008), and cortex (009, 010, 011). (**B**) Analytic scheme for DEGs analysis. First, a global comparison of AD-related DEGs versus control at early- (light grey) and late-stage (dark grey) disease was performed in all neurons; genes unique to the non-neuronal background were excluded. Second, DEGs unique to early- or late-stage disease were sub-analyzed by interneuron subtype (Pv yellow color; green color)- and excitatory neuron (red color)-specific gene expression. Finally, regional specificity by DG (dashed), CA1 (bold), CA3 (dotted), and cortex (double-line) of Pv- or Sst-specific interneuron DEGs was analyzed.

Segmented areas were exposed to automatically controlled UV laser to cleave probe barcodes from specific cell regions within ROIs. Finally, a “background” region of all unstained areas/cells within the ROI was collected. Liberated barcodes from each ROI were collected by microcapillary and plated into individual wells of a microtiter plate before processing by the NanoString Max/Flex nCounter system.

### Counting GeoMx photocleaved oligos by nCounter hybridization assay

Probes cleaved from the GeoMx DSP spatial analysis of brain ROIs were counted using nCounter readout, according to manufacturer instructions (NanoString, manual MAN-10089-08). Briefly, samples of GeoMx DSP-cleaved probes were aspirated dry, rehydrated in distilled, nuclease-free water, and transferred to another plate where they were incubated overnight (16 to 24 h) with hybridization probes (Hyb Code Pack, NanoString) in a thermal cycler set to 65 °C with heated lid (70 °C). Each Hyb Code hybridization probe contains a barcoded reporter tag to identify sequences in target GeoMx DSP-cleaved probes and a capture tag to collect hybridized complexes. Samples were then pooled by column into 12-well strip tube and processed using a NanoString MAX/FLEX/Pro Prep Station in high sensitivity mode. Data were acquired using a NanoString Digital Analyzer with 555 fields of view setting.

### Data and image analysis for spatial transcriptomics

Data analysis was performed using NanoString-developed software (GeoMxTools 3.5.0^26^ and NanoStringNCTools^27^) combined with open-source R package software. In the overall process, raw gene expression counts were subjected to dimension reduction and differential gene expression analysis, which were visually explored. Analyses were performed separately on two datasets, gene counts from early-stage as well as late-stage brain ROIs, to avoid potential batch effects.

First, Digital Count Conversion (DCC) files, containing expression count data and sequencing quality metadata, and Probe Kit Configuration (PKC) files, containing probe assay and sample metadata, were imported into R. Gene expression counts with a value of 0 were shifted to 1 to facilitate downstream transformations. ROI segments were filtered and removed if they met the criteria: “<1000 raw reads”, “below 80% Aligned or 80% Trimmed”, “80% Stitched”, “below 50% sequencing saturation”. Genes were filtered and removed if they were not detected in at least 10% of segments. Next, counts were normalized using quartile 3 normalization and visualized by uniform manifold approximation and projection, a non-linear dimensionality reduction visualization tool. Gene expression in both datasets was relatively well separated according to ROI brain region.

Finally, differential expression was performed by linear mixed-effect model (LMM). LMM accounts for subsampling across a tissue or a slide by adjusting for multiple ROIs, which are not independent observations, within each tissue section. This contrasts with traditional statistical tests, which are only valid for independent observations. Formulating the LMM model depends on the scientific query. The LMM model does not require a random slope if comparing mutually exclusive features, *e.g*., 5XFAD in CA1 versus control in CA1. The LMM model selected was log_2_(gene)∼genotype+(1|slide). Including the “slide” as a random variable in the LMM model accounted for inter-slide variation in identifying DEGs. DEGs were identified using the GeoMx analysis suite, which utilizes the GeoMxTools R package (“mixedModelDE” in R package “lmerTest”). Overlap between DEGs in different regions were identified using VennDiagram R package.

## RESULTS

### Spatial transcriptomics of mouse brain and identification of interneurons

Herein, we employed the 5XFAD mouse model of AD to investigate interneuron dysfunction spatially and temporally with disease progress. The 5XFAD mouse develops brain amyloid plaques, neurodegeneration, microgliosis, and cognitive impairment mirroring humans. The 5XFAD mouse also develops interneuron dysfunction, with neural circuit overactivity^20, 22^ and GABA inhibitory interneuron loss within the hippocampus,^28^ also recapitulating features of AD in humans. Spatial transcriptomics of brain sections from 5XFAD versus non-carrier littermates was leveraged to evaluate interneuron dysfunction at the early (12 wk of age) and late (30 wk of age) stages of AD. Interneurons were identified by Pv and Sst staining and excitatory neurons by vesicular glutamate transporter (Vglut) staining by FISH of brain sections, which were then analyzed by spatial transcriptomics (**Figure 1A**). Differentially expressed genes (DEGs) were identified within regions of interest (ROIs) and used to examine biological differences in 5XFAD versus control mice at early- and late-stage disease (**Figure 1B**). DEGs were then also analyzed in 5XFAD versus control mice by interneuron- (PV, SST) and excitatory neuron-specificity (**Figure 1C**) and also by brain region (dentate gyrus, hippocampal CA1 and CA3, and cortex, **Figure 1D**).

### Global DEGs in AD versus control neurons at early- and late-stage disease

The total number of DEGs differentiating 5XFAD from control animals increased from 9,306 at early-stage to 18,000 at late-stage disease, as might be anticipated with worsening illness. Overview analysis in 5XFAD versus control mice identified 1,318 shared DEGs and 7,988 unique DEGs at early-stage and 16,682 unique DEGs at late-stage disease, after subtraction of all genes not confined to Pv-, Sst-, and Vglut-positive neurons (**Figure 2A**; **Supplementary Table S1**). Pathway enrichment analysis of DEGs shared between early- and late-stage using Kyoto Encyclopedia of Genes and Genomes (KEGG) terms found “pathways of neurodegeneration − multiple diseases” was most significant and with the most DEG members in 5XFAD brain, closely followed with “Parkinson disease”, “Huntington disease”, “Alzheimer’s disease”, “amyotrophic lateral sclerosis”, and “prion disease” (**Figure 2B**; **Supplementary Table S2** for KEGG; **Supplementary Table S3** for Gene Ontology [GO]). These pathways were also represented in DEGs unique to early-stage disease, but to a less significant extent, albeit with more DEG members (**Figure 2C**; **Supplementary Tables S2-S3**). Various biological pathways related to RNA and protein processing, transport, and clearance, such as “spliceosome”, “protein processing in endoplasmic reticulum”, “nucleocytoplasmic transport”, and “autophagy - animal” were highly significant in early-stage 5XFAD versus control brain. At late-stage, pathways related to neurodegeneration grew more significant with more DEG members, aligned with AD progression (**Figure 2D**; **Supplementary Tables S2-S3**).

**Figure 2.**
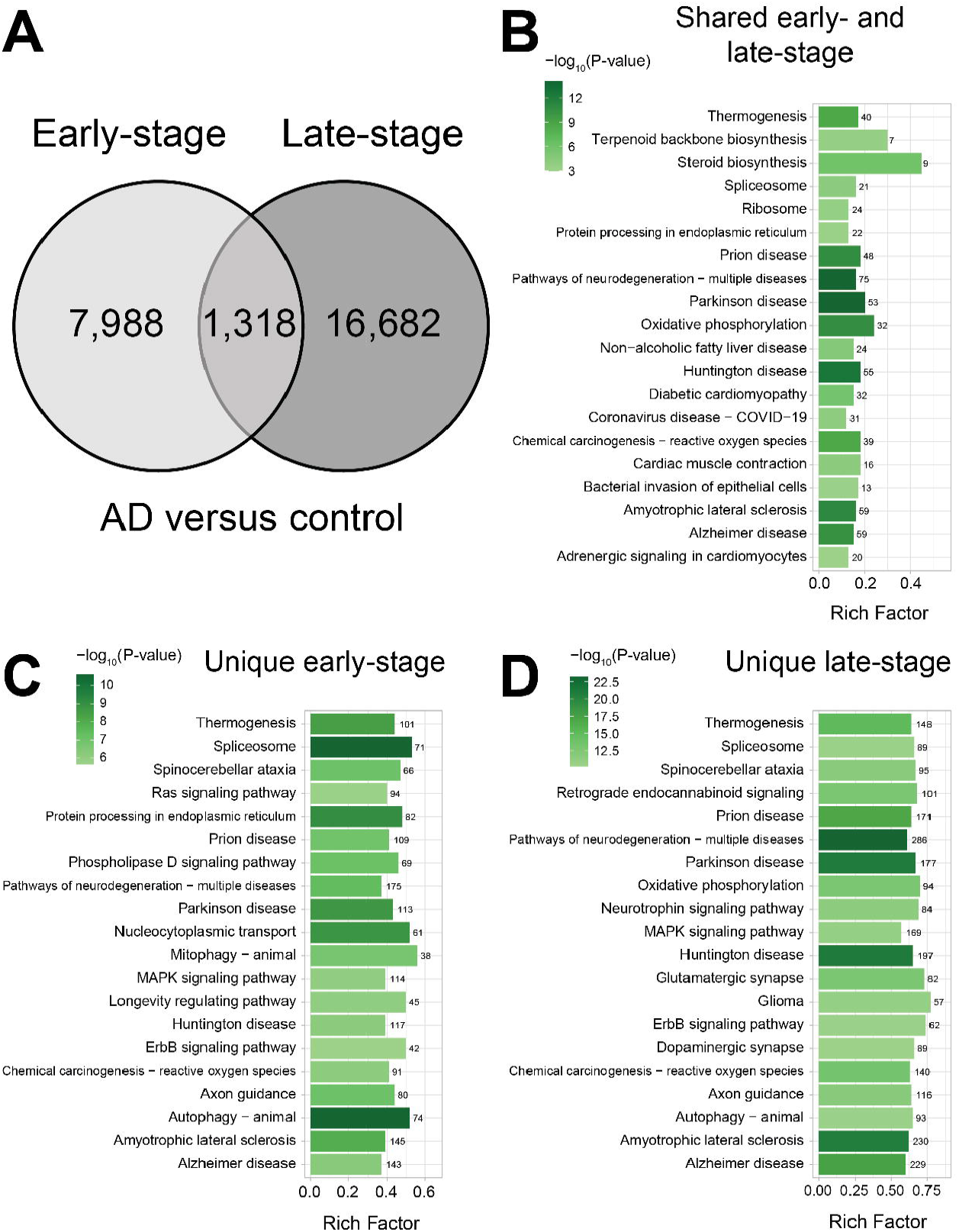
Global DEGs in AD versus control brain at early- and late-stage disease. (**A**) Venn diagram of differentially expressed genes (DEGs) in 5XFAD AD versus control brain at early-stage (light grey) and late-stage (dark grey) disease. KEGG pathway enrichment analysis of DEGs (**B**) shared between early- and late-stage disease, (**C**) unique to early-stage disease, and (**D**) unique to late-stage disease. Legend represents significance level by -log_10_(p-value), numbers over the bars represent number of DEGs belonging to annotated pathways. Background non-neuronal DEGs were subtracted from the analysis.

### Interneuron- and excitatory neuron-specific DEGs in AD versus control at early- and late-stage disease

Following the global overview analysis, we next homed in on interneuron- and excitatory neuron-specific DEGs in 5XFAD versus control brain at early- and late-stage disease. In early-stage disease, all neurons shared 11 DEGs and Pv- and Sst-positive interneurons shared 94 DEGs (**Figure 3A**; **Supplementary Table S4**). Interestingly, excitatory neurons shared more DEGs with either interneuron subtype, 149 with Pv-positive and 178 with Sst-positive, than the interneurons shared with each other. Most DEGs were uniquely expressed by various neuron types, with 1,562 DEGs uniquely expressed by Pv-positive interneurons, 2,284 DEGs uniquely expressed by Sst-positive interneurons, and 3,267 DEGs uniquely expressed by Vglut-positive excitatory neurons. DEGs were only nominally significant in interneurons, compared to excitatory neurons, which expressed DEGs that remained significant after multiple adjustments. There are far fewer interneurons than excitatory neurons in the brain^29^ and in our ROIs (**Figure 1**), lowering the resolution to detect DEGs in interneurons.

**Figure 3.**
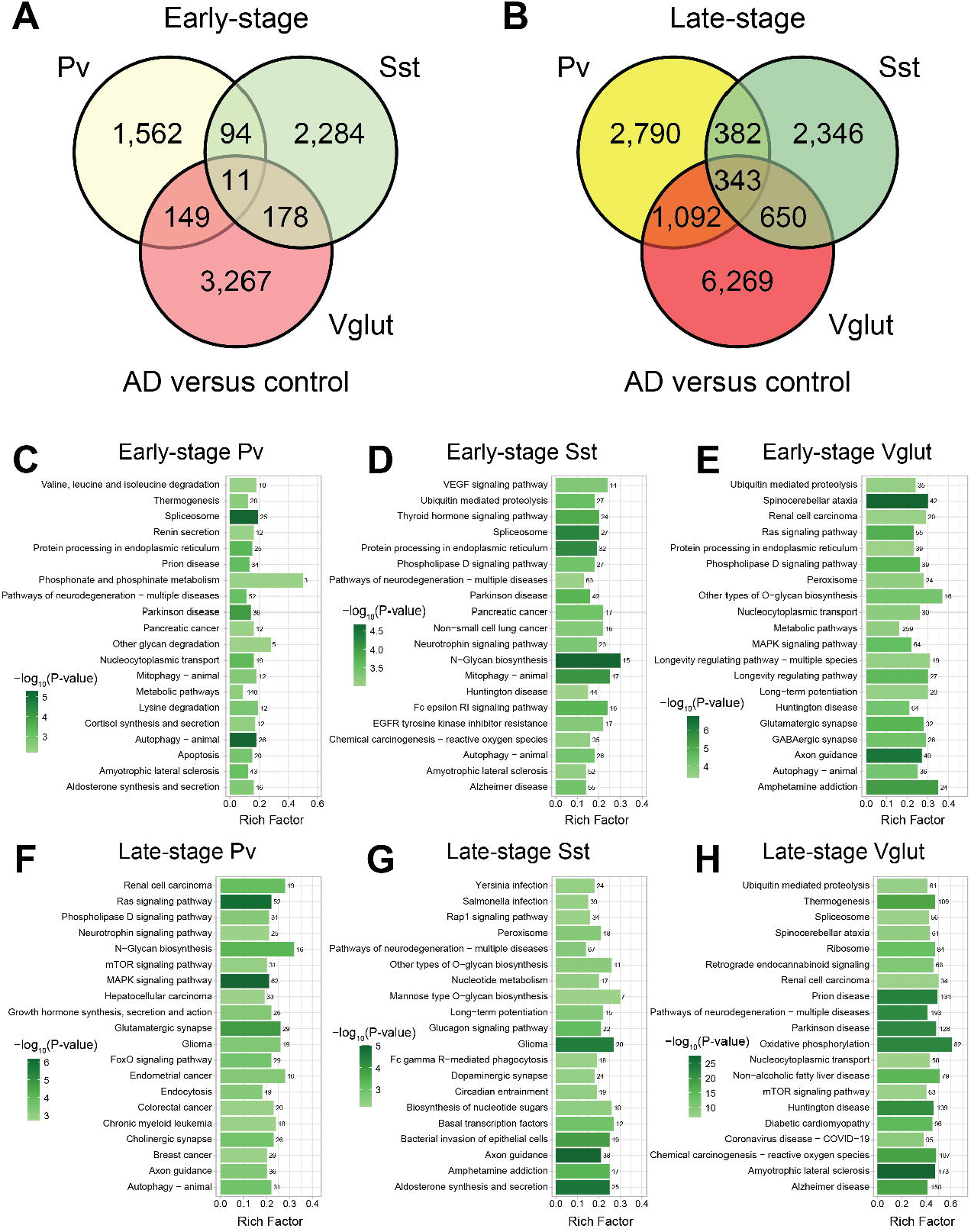
Interneuron- and excitatory neuron-specific DEGs in AD versus control at early- and late-stage disease. Venn diagram of differentially expressed genes (DEGs) in 5XFAD AD versus control across Pv- (yellow) and Sst (green)-positive interneurons and Vglut (red)-positive excitatory neurons at (**A**) early-stage (light shading) and (**B**) late-stage (dark shading) disease. KEGG pathway enrichment analysis of early-stage DEGs in 5XFAD AD versus control in (**C**) Pv- positive interneurons, (**D**) Sst-positive interneurons, and (**E**) Vglut-positive excitatory neurons. KEGG pathway enrichment analysis of late-stage DEGs in 5XFAD AD versus control in (**F**) Pv- positive interneurons, (**G**) Sst-positive interneurons, and (**H**) Vglut-positive excitatory neurons. Legend represents significance level by -log_10_(p-value), numbers over the bars represent number of DEGs belonging to annotated pathways. Background non-neuronal DEGs were subtracted from the analysis.

KEGG pathway enrichment of unique DEGs in Pv-positive interneurons at early-stage yielded candidates connected to RNA and protein metabolism, such as “autophagy – animal”, “spliceosome”, “protein processing in endoplasmic reticulum”, and “nucleocytoplasmic transport”, as well as neurodegeneration, *e.g*., “Parkinson disease”, “prion disease” *etc.* (**Figure 3C**; **Supplementary Table S5** for KEGG; **Supplementary Table S6** for GO). “Metabolic pathways” contained the most DEGs by far, at 140. Pathway analysis of Sst-positive interneurons at early-stage found a distinct biology; while several hits prominent in Pv-positive interneurons remained, such as “spliceosome” and “protein processing in endoplasmic reticulum”, novel hits emerged in Sst-positive interneurons, such as “N-glycan biosynthesis” and “thyroid hormone signaling pathway” (**Figure 3D**; **Supplementary Tables S5-S6**). Finally, excitatory neurons were distinct from interneurons; neurodegenerative pathways were present, but “spinocerebellar ataxia” was the most significant (**Figure 3E**; **Supplementary Tables S5-S6**). “Axon guidance” was also highly characteristic of excitatory neurons, along with synapse-related pathways, including “glutamatergic synapse” and, surprisingly, “GABAergic synapse”.

At late-stage disease, all neurons shared 343 DEGs and Pv- and Sst-positive interneurons shared 382 DEGs (**Figure 3B**). Like the early time point, excitatory neurons shared more DEGs with either interneuron subtype, 1,092 with Pv-positive and 650 with Sst-positive, than the interneurons shared with each other. Moreover, there were more shared and unique DEGs at late-stage than at early-stage, possibly indicative of more progressive disease, which leads to greater transcriptomics differences in 5XFAD versus control neurons. Also, similar to the early stage, most DEGs were uniquely expressed, with 2,790 DEGs expressed by Pv-positive interneurons, 2,346 DEGs uniquely expressed by Sst-positive interneurons, and 6,269 DEGs uniquely expressed by Vglut-positive excitatory neurons (**Supplementary Table S7**).

KEGG pathway enrichment of DEGs from late-stage Pv-positive interneurons suggested predominance of conventionally oncogenic signaling pathways, including “MAPK signaling pathway” and “Ras signaling pathway” as the most and second most significant containing the most DEGs (**Figure 3F**; **Supplementary Table S8** for KEGG; **Supplementary Table S9** for GO). These were accompanied by several cancer pathways, *e.g*., renal cell carcinoma and glioma. Neuronal pathways were also represented, such as “axon guidance” and “neurotrophin signaling pathway”. Interestingly, excitatory, *i.e*., “glutamatergic synapse”, and neuromodulatory, *i.e*., “cholinergic synapse” neurotransmission pathways were present. Within Sst-positive interneurons, “axon guidance” was the most significant pathways, followed by “aldosterone synthesis and secretion” and “glioma” (**Figure 3G**; **Supplementary Tables S8-S9**). Glycan biosynthesis was represented and was more significant than in Pv-interneurons. Moreover, several pathways related to infection and clearance, “bacterial invasion of epithelial cells”, “Yersinia infection”, “Salmonella infection”, and “Fc gamma R-mediated phagocytosis”. In contrast to interneurons, late-stage excitatory neurons continued to be characterized by pathways related to neurodegeneration, top among them “amyotrophic lateral sclerosis”, but also “prion disease”, “Parkinson disease”, “pathways of neurodegeneration - multiple diseases”, “Huntington disease”, and “Alzheimer disease” (**Figure 3H**; **Supplementary Tables S8-S9**).

### Region-specific DEGs in AD versus control Pv- and Sst-positive interneurons at early- and late-stage disease

Next, we performed a DEGs analysis in 5XFAD versus control by spatial location, given the heterogeneity of the brain as well as the spatial-dependent spread in pathology and neuronal loss. At early-stage in Pv-positive interneurons, there were no DEGs shared across all examined regions, the dentate gyrus (DG), CA1, CA3, and cortex (**Figure 4A**; **Supplementary Table S10**). Generally, few DEGs overlapped between 2 or 3 regions overall. Early-stage Pv-positive interneurons in the DG uniquely expressed 336 DEGs, which KEGG pathway analysis showed were related most significantly to “autophagy – animal” and pathways containing the most DEG, 10, were “Ras signaling” and “coronavirus disease – COVID–19” pathways (**Figure 4B**; **Supplementary Table S11** for KEGG; **Supplementary Table S12** for GO). Early-stage Pv-positive interneurons in CA1 uniquely expressed 310 DEGs that differed in 5XFAD versus control, correlated most significantly to the “spliceosome” pathway with the “metabolic pathways” having the most DEGs, 34 (**Figure 4C**). Early-stage Pv-positive interneurons in CA3 uniquely expressed 333 DEGs, linked most significantly to “chemical carcinogenesis – reactive oxygen species” and “metabolic pathways” again encompassing the most DEGs, 42 (**Figure 4D**). Early-stage Pv-positive interneurons residing in the cortex, the last examined brain region, uniquely expressed 413 DEGs, related most significantly to “pathways in cancer”, which also contained the most DEGs, 21 (**Figure 4E**).

**Figure 4.**
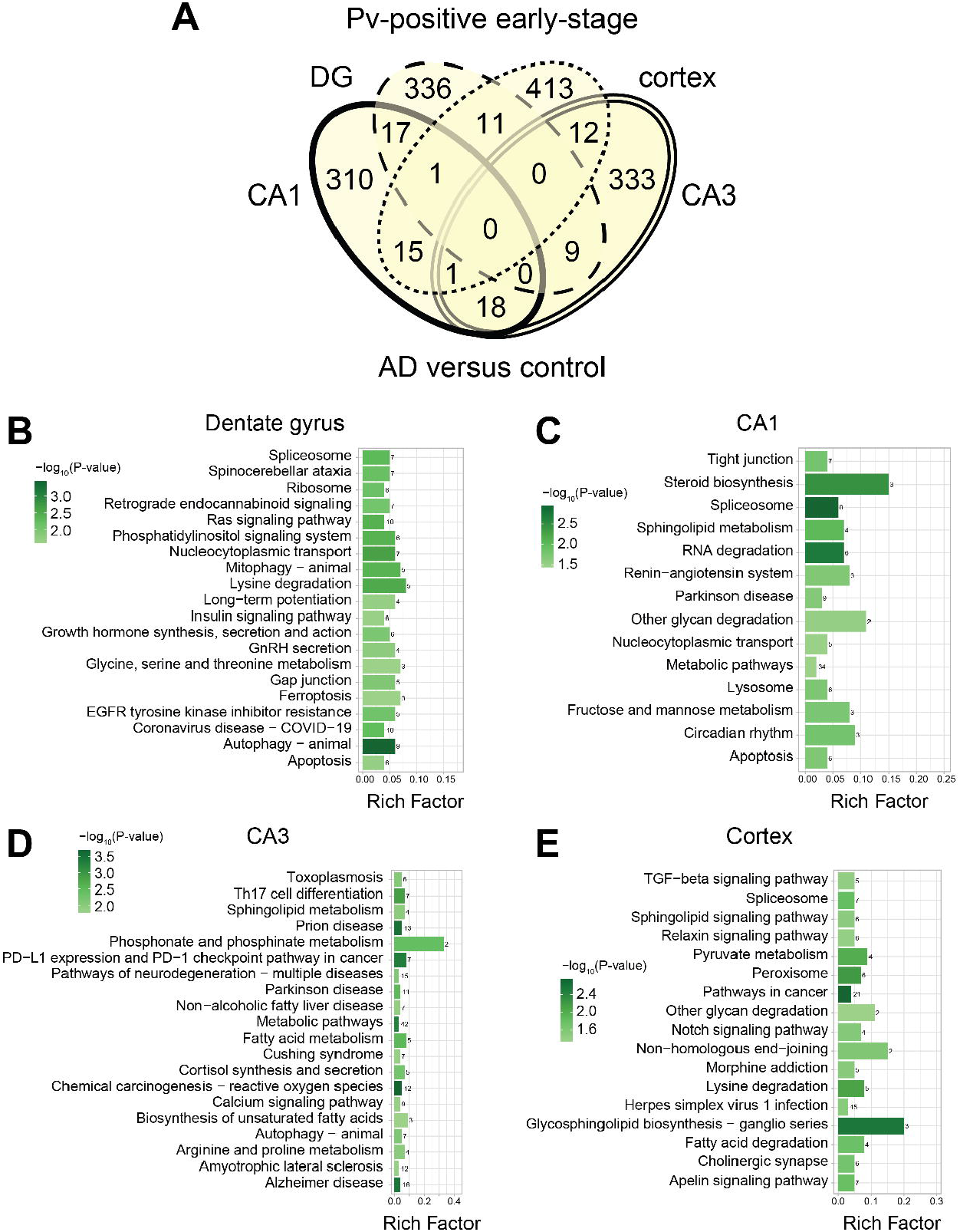
Region-specific DEGs in AD versus control Pv-positive interneurons at early-stage disease. (**A**) Venn diagram of differentially expressed genes (DEGs) in 5XFAD AD versus control across all analyzed regions dentate gyrus (DG), CA1, CA3, and cortex in Pv-positive interneurons at early-stage disease. KEGG pathway enrichment analysis of early-stage DEGs in 5XFAD AD versus control Pv-positive interneurons in (**B**) DG (dashed), (**C**) CA1 (bold), (**D**) CA3 (dotted), (**E**) cortex (double-line). Legend represents significance level by -log_10_(p-value), numbers over the bars represent number of DEGs belonging to annotated pathways. Background non-neuronal DEGs were subtracted from the analysis.

During early-stage disease, in Sst-positive interneurons, there were no 5XFAD versus control DEGs shared by all regions DG, CA1, CA3, and cortex, and proportionately few by 2 or 3 regions (**Figure 5A**; **Supplementary Table S13**). Early-stage Sst-positive interneurons in the DG uniquely expressed 671 DEGs, related most significantly to “terpenoid backbone biosynthesis”, and with “metabolic pathways” containing the highest number of DEGs, 63 (**Figure 5B**; **Supplementary Table S14** for KEGG; **Supplementary Table S15** for GO). Early-stage Sst-positive interneurons in CA1 uniquely expressed 422 DEGs, which pathway analysis found was most significantly linked to “chronic myeloid leukemia” (**Figure 5C**) and “amyotrophic lateral sclerosis” and “Alzheimer disease” contained the most DEGs, 14. Early-stage Sst-positive interneurons in CA3 uniquely expressed 348 DEGs, related most significantly to “N-glycan biosynthesis” and with “spliceosome” encompassing the most DEGs, 9 (**Figure 5D**). Early-stage Sst-positive interneurons located in the cortex uniquely expressed 488 DEGs, related most significantly to the “ErbB signaling pathway” and with “pathways of neurodegeneration - multiple diseases” containing the largest number of DEGs, 16 (**Figure 5E**).

**Figure 5.**
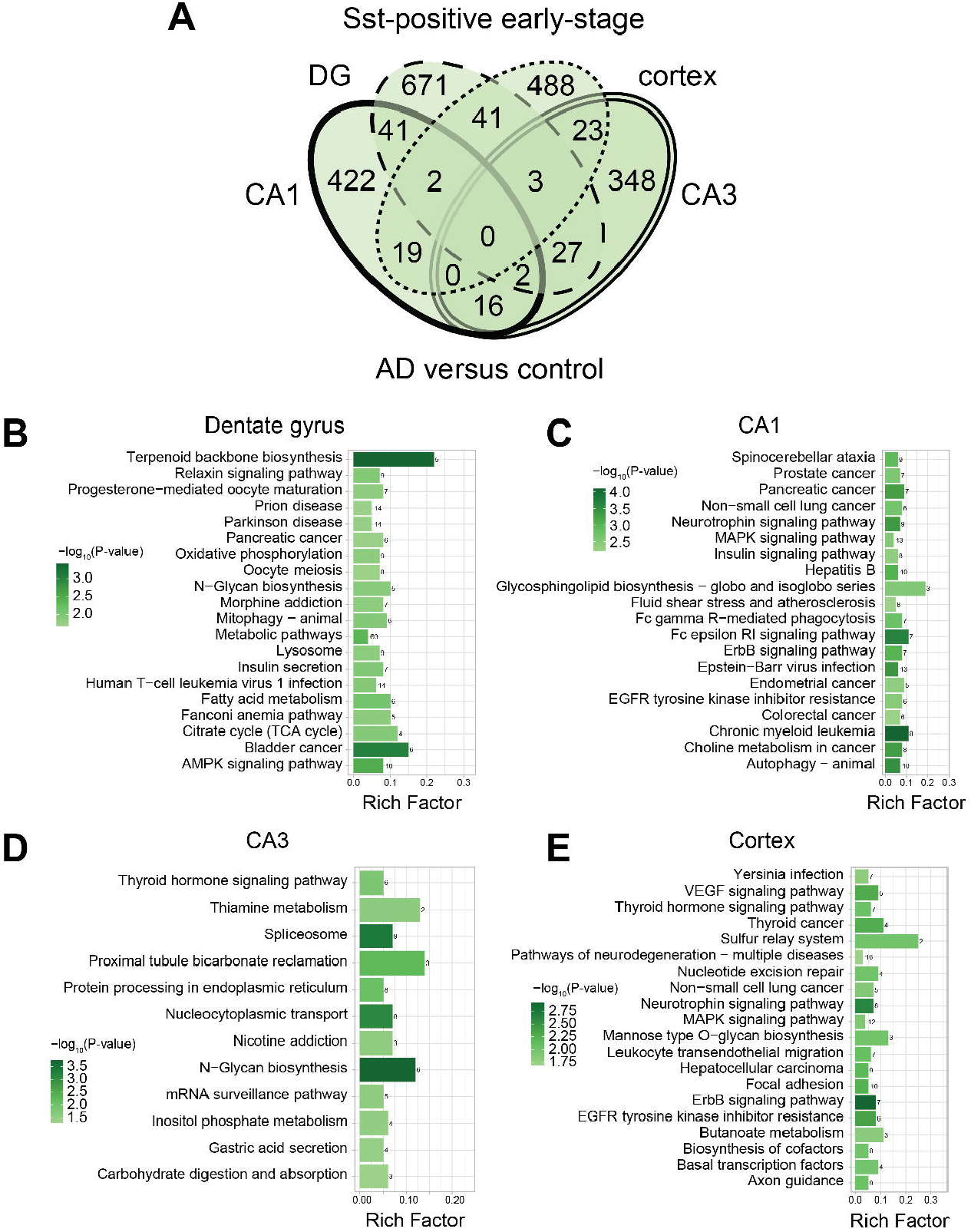
Region-specific DEGs in AD versus control Sst-positive interneurons at early-stage disease. (**A**) Venn diagram of differentially expressed genes (DEGs) in 5XFAD AD versus control across all analyzed regions dentate gyrus (DG), CA1, CA3, and cortex in Sst-positive interneurons at early-stage disease. KEGG pathway enrichment analysis of early-stage DEGs in 5XFAD AD versus control Sst-positive interneurons in (**B**) DG (dashed), (**C**) CA1 (bold), (**D**) CA3 (dotted), (**E**) cortex (double-line). Legend represents significance level by -log_10_(p-value), numbers over the bars represent number of DEGs belonging to annotated pathways. Background non-neuronal DEGs were subtracted from the analysis.

When we examined DEGs in early-stage 5XFAD versus control excitatory Vglut-positive neurons, again, none overlapped across all brain regions, and relatively few were shared between 2 or 3 regions (**Supplementary Figure S1A**; **Supplementary Table S16**). However, there were generally more DEGs in excitatory neurons compared to inhibitory interneurons, as anticipated. Early-stage Vglut-positive excitatory neurons in the DG uniquely expressed 868 DEGs, related most significantly to “glutamatergic synapse”, with “axon guidance”, the second most significant, having the most DEGs, 19 (**Supplementary Figure S1B**; **Supplementary Table S17** for KEGG; **Supplementary Table S18** for GO). Early-stage Vglut-positive excitatory neurons in CA1 uniquely expressed 878 DEGs, correlated most significantly to “spinocerebellar ataxia” (**Supplementary Figure S1C**) and with “Ras signaling pathway”, “calcium signaling pathway”, and “endocytosis” encompassing the most DEGs, 19. Early-stage Vglut-positive excitatory neurons in CA3 uniquely expressed 217 DEGs, linked most significantly to “small cell lung cancer” and “metabolic pathways” having the most DEGs, 21 (**Supplementary Figure S1D**). Early-stage Vglut-positive excitatory neurons located in the cortex uniquely expressed 570 DEGs, related most significantly to “thermogenesis” and “metabolic pathways”, the second most significant, comprising the largest number of DEGs, 69 (**Supplementary Figure S1E**).

Finally, we examined DEGs and pathways in late-stage 5XFAD versus control in neurons by brain region. At late-stage in Pv-positive interneurons, 2 DEGs were shared across all examined regions, DG, CA1, CA3, and cortex, although, generally, proportionality few DEGs overlapped between 2 or 3 regions (**Figure 6A**; **Supplementary Table S19**). Late-stage Pv-positive interneurons in the DG uniquely expressed 460 DEGs, which pathway analysis found was most significantly related to the “viral carcinogenesis”, which also contained the most DEG members, 11 (**Figure 6B**; **Supplementary Table S20** for KEGG; **Supplementary Table S21** for GO). Late-stage Pv-positive interneurons in CA1 uniquely expressed 331 DEGs, related most significantly to the “mTOR signaling pathway” pathway with “metabolic pathways” having the most DEGs, at 34 (**Figure 6C**). Late-stage Pv-positive interneurons in CA3 uniquely expressed 833 DEGs, enriched most significantly in “MAPK signaling pathway”, which also had the most DEGs, 27 (**Figure 6D**). Late-stage Pv-positive interneurons located in the cortex uniquely expressed 659 DEGs, related most significantly to “pathways in cancer”, which also comprised the most DEGs, 25 (**Figure 6E**).

**Figure 6.**
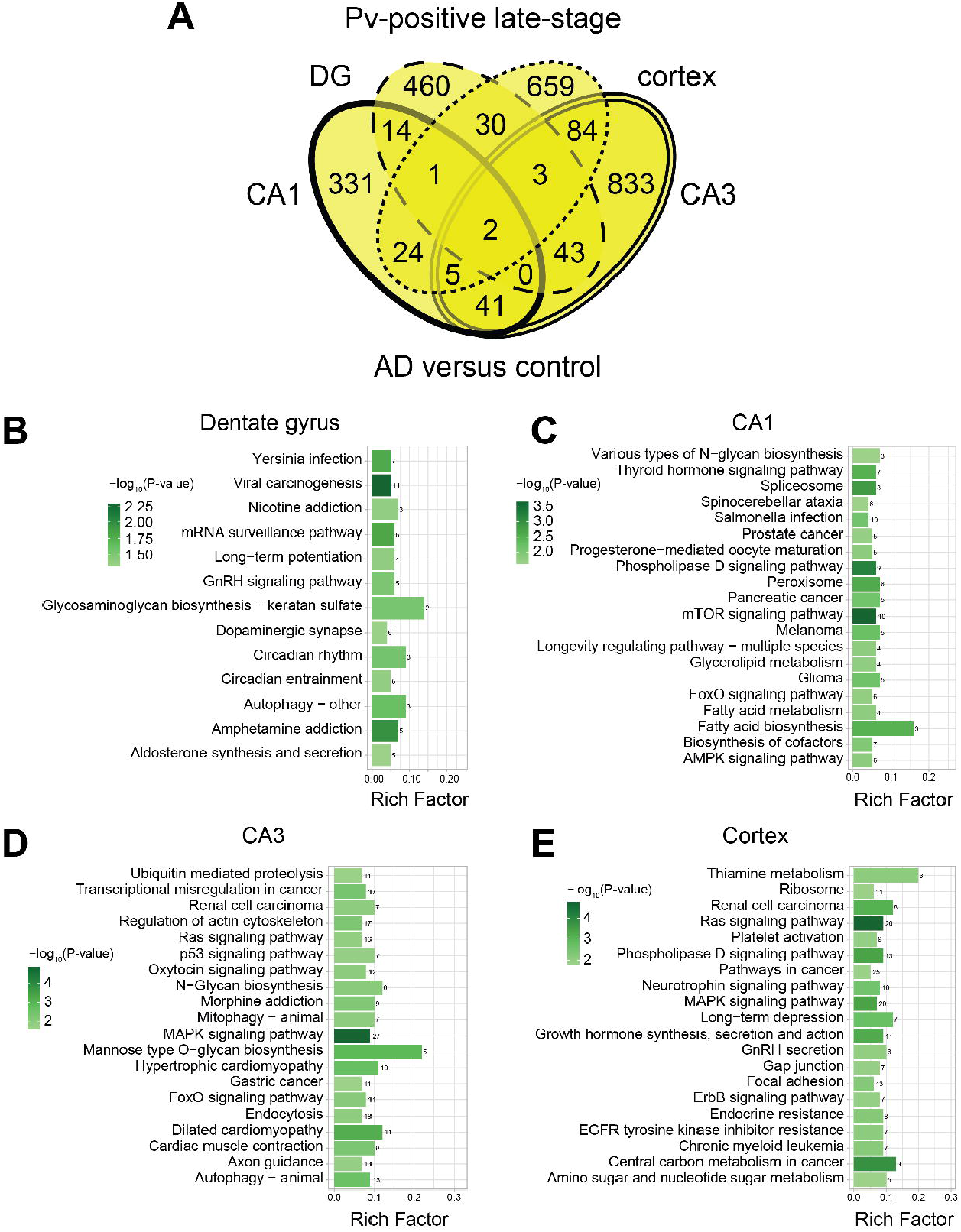
Region-specific DEGs in AD versus control Pv-positive interneurons at late-stage disease. (**A**) Venn diagram of differentially expressed genes (DEGs) in 5XFAD AD versus control across all analyzed regions dentate gyrus (DG), CA1, CA3, and cortex in Pv-positive interneurons at late-stage disease. KEGG pathway enrichment analysis of late-stage DEGs in 5XFAD AD versus control Pv-positive interneurons in (**B**) DG (dashed), (**C**) CA1 (bold), (**D**) CA3 (dotted), (**E**) cortex (double-line). Legend represents significance level by -log_10_(p-value), numbers over the bars represent number of DEGs belonging to annotated pathways. Background non-neuronal DEGs were subtracted from the analysis.

At late-stage disease, in Sst-positive interneurons, there were no 5XFAD versus control DEGs shared by all regions, and comparatively scarce DEGs overlapping between 2 or 3 regions (**Figure 7A**; **Supplementary Table S22**). Late-stage Sst-positive interneurons in the DG uniquely expressed 464 DEGs, related most significantly to “axon guidance”, which also contained most DEGs, 11 (**Figure 7B**; **Supplementary Table S23** for KEGG; **Supplementary Table S24** for GO). Late-stage Sst-positive interneurons in CA1 uniquely expressed 428 DEGs, which pathway analysis found was most significantly linked to “Parkinson disease” (**Figure 7C**) with “pathways of neurodegeneration - multiple diseases” having the highest number of DEGs, 20. Late-stage Sst-positive interneurons in CA3 uniquely expressed 629 DEGs, related most significantly to “bacterial invasion of epithelial cells” and with “metabolic pathways” comprising the most DEGs, 58 (**Figure 7D**). Late-stage Sst-positive interneurons within uniquely expressed 460 DEGs, related most significantly to “carbon metabolism” along with other metabolic “oxidative phosphorylation” and “metabolic pathways” containing the highest number of DEGs, 50 (**Figure 7E**).

**Figure 7.**
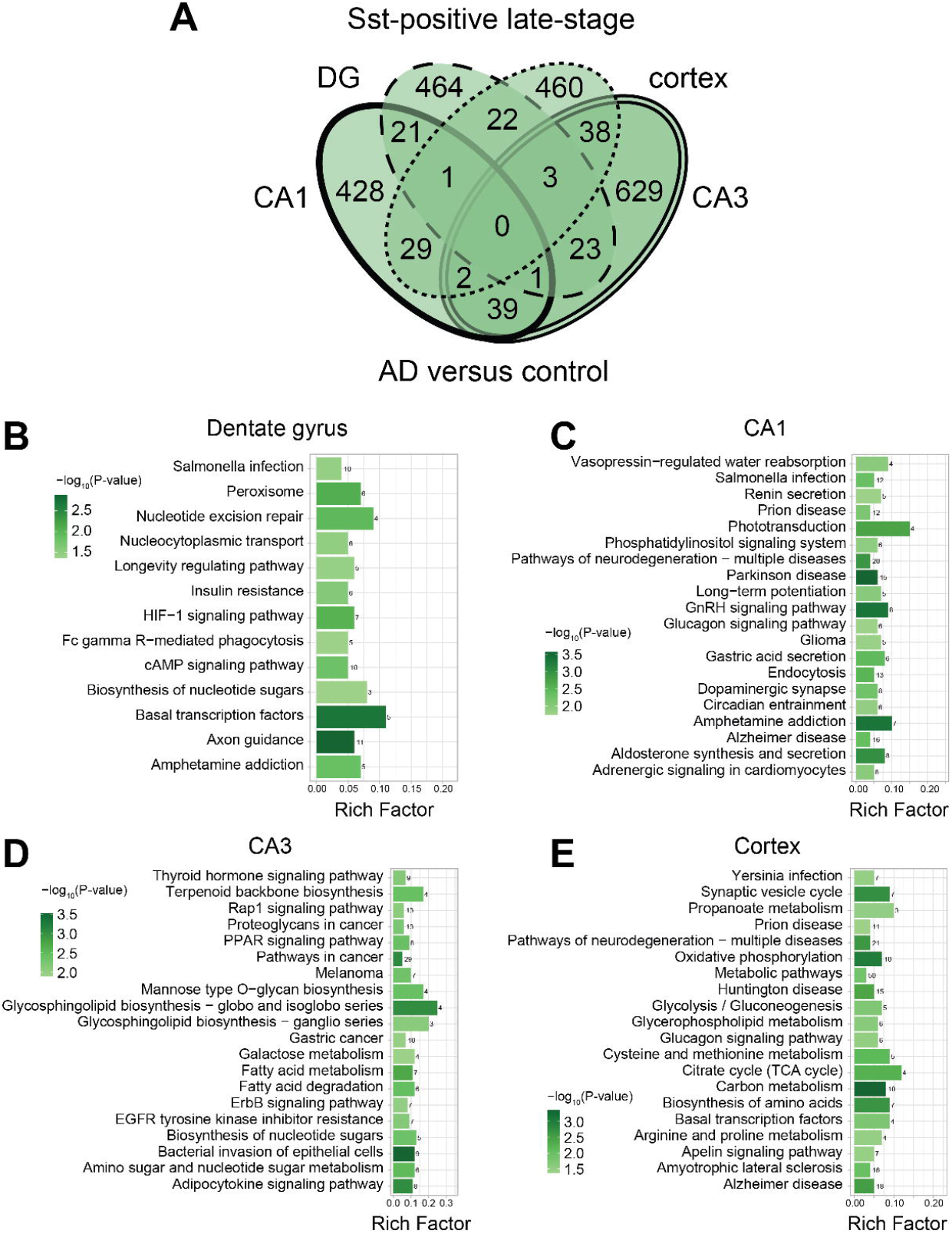
Region-specific DEGs in AD versus control Sst-positive interneurons at late-stage disease. (**A**) Venn diagram of differentially expressed genes (DEGs) in 5XFAD AD versus control across all analyzed regions dentate gyrus (DG), CA1, CA3, and cortex in Sst-positive interneurons at late-stage disease. KEGG pathway enrichment analysis of late-stage DEGs in 5XFAD AD versus control Sst-positive interneurons in (**B**) DG (dashed), (**C**) CA1 (bold), (**D**) CA3 (dotted), (**E**) cortex (double-line). Legend represents significance level by -log_10_(p-value), numbers over the bars represent number of DEGs belonging to annotated pathways. Background non-neuronal DEGs were subtracted from the analysis.

When we assessed DEGs in late-stage 5XFAD versus control excitatory Vglut-positive neurons, 51 were shared by all brain regions along with several overlapping between 2 or 3 various regions (**Supplementary Figure S2A**; **Supplementary Table S25**). Late-stage Vglut-positive excitatory neurons in the DG uniquely expressed 810 DEGs, related most significantly to “synaptic vesicle cycle” with “thermogenesis”, the second most significant, containing most DEGs, 26 (**Supplementary Figure S2B**; **Supplementary Table S26** for KEGG; **Supplementary Table S27** for GO). Late-stage Vglut-positive excitatory neurons in CA1 uniquely expressed 433 DEGs, correlated most significantly to “lysosome” with “metabolic pathways” encompassing the most DEGs, 46 (**Supplementary Figure S2C**). Late-stage Vglut-positive excitatory neurons in CA3 uniquely expressed 1,548 DEGs, linked most significantly to “amyotrophic lateral sclerosis” and other neurodegenerative pathways with “metabolic pathways” having the most DEGs, 122 (**Supplementary Figure S2D**). Late-stage cortical Vglut-positive excitatory neurons uniquely expressed 579 DEGs, related most significantly to “sphingolipid signaling pathway” and “pathways in cancer” comprising the most DEGs, 23 (**Supplementary Figure S2E**).

## DISCUSSION

Imbalance occurs in neuronal excitation versus inhibition in the AD brain, leading to network hyperexcitability.^7–9^ Impaired interneuron activity also curbs neurogenesis, GABAergic transmission,^9–11^ dynamics of Aβ clearance,^9, 11, 20, 22–24^ and cognitive functioning^11^ with ultimate loss of interneurons in both humans and mouse models.^9, 11, 30^ Critically, as a potential therapeutic target, interneuron dysfunction may be an early pathological event preceding clinical manifestations of AD.^12–17^ This is borne out by preclinical studies that suggest feasibility and salutary effects from interneuron modulation;^19, 20^ however, deeper understanding of interneuron biology in the context of AD is needed before they can be leveraged to develop mechanism-based and targeted therapies.

To fill this knowledge gap, we launched the present study to generate a roadmap of interneuron dysfunction in AD by profiling interneuron-specific gene expression in 5XFAD AD versus control brain. Our study comprised two time points; early-stage and presymptomatic to disease in mice aged 12 weeks to examine potentially initiating events contributing to pathology, which could constitute prime targets for intervention before symptom onset. We also assessed gene expression at late-stage disease in mice aged 30 weeks that may serve as markers of disease progression and severity. We leveraged a spatial platform of gene expression analysis due to the heterogeneity of the brain, which could also impact anatomic heterogeneity of interneuron function and pathology. Moreover, Aβ spreads across various brain areas with disease progression in humans^31^ and 5XFAD mice,^30, 32, 33^ and deposits may be linked to interneuron activity;^20^ therefore, spatial analysis accounts for this heterogeneity.

We analyzed DEGs from our spatial dataset followed by pathway enrichment analysis at three levels, global (early-versus late-stage), neuron-specific (Pv-, Sst-, and excitatory-specific), and region-dependent (DG, CA1, CA3, cortex). The global analysis revealed changes overall in 5XFAD versus control brain with disease progression. Various biological pathways related to RNA and protein processing, transport, and clearance were hallmarks of early-stage AD, with top significant hits such as “spliceosome”, “protein processing in endoplasmic reticulum”, “nucleocytoplasmic transport”, and “autophagy - animal”, as well as less significant pathways, *e.g*., “lysosome”. Moreover, rich factors were relatively high for these pathways, around 0.5, suggesting that half of all DEG members belonging to these KEGG pathways were represented in our dataset. Defects in RNA^34–38^ and especially protein^39–42^ metabolism and/or transport are widely reported in AD; pharmacologically or genetically modulating the activity of these biological processes indicate they may drive or significantly contribute to disease progression. Pathway analysis also enriched “mitophagy - animal”, another early and key attribute of AD.^43^

In addition to these biological processes, neurodegenerative pathways were also present at early-stage disease, including “pathways of neurodegeneration − multiple diseases”, “Parkinson disease”, “Huntington disease”, “Alzheimer’s disease”, “amyotrophic lateral sclerosis”, and “prion disease”, which were the most significant pathways shared with late-stage disease. Importantly, rich factors for neurodegenerative pathways were around 0.4 at early-stage AD, but increased to around 0.6 at late-stage, implying greater DEGs representation from these pathways in our dataset, as would be anticipated with disease progression. Impaired RNA and protein metabolism continued to be represented at late-stage, but were less significant than neurodegenerative pathways, the inverse to the early-stage profile.

Next, we homed in on neuron-specific changes occurring in AD versus control at early- and late-stage disease, examining Pv- and Sst-positive interneurons as well as Vglut-positive excitatory neurons. Early stage Pv-positive interneurons reflected the same pathways that emerged from the global analysis, with compromised RNA and protein metabolism and transport as the most salient pathways, followed closely by neurodegenerative pathways. Sst-positive interneurons exhibited a similar profile but there were fewer neurodegenerative pathways in the tip 10 hits. Moreover, a salient difference was the most significant pathway, which was “N-glycan biosynthesis”, related to endoplasmic reticulum processing of glycans on protein surfaces, which also had the highest rich factor. There is a small but emerging literature regarding altered N-glycosylation in AD,^44–46^ although the significance remains incompletely understood.^47^

Interestingly, excitatory Vglut-positive neurons had a distinct profile from interneurons; only the neurodegenerative pathway “spinocerebellar ataxia” was present among the top 10 significant pathways. Based on the stronger presence of early-stage neurodegenerative markers in Pv-positive interneurons, it is tempting to speculate that they may be affected by AD before excitatory neurons and constitute an early event in the disease course. Instead, excitatory neurons were characterized by several axon- and synapse-related pathways, such as “axon guidance”, “glutamatergic synapse” and, surprisingly, “GABAergic synapse”. “Amphetamine addiction” was also featured, related to dopamine levels and long-term adaptive brain changes. “Longevity regulating pathway”, primarily metabolic, and “Ras signaling pathway”, “phospholipase D signaling pathway”, and “MAPK signaling pathway”, principally cell-regulating, were also present. Ras homolog gene family member A (RhoA), a DEG present in our dataset and belong to “Ras signaling pathway”, is a small GTPase belonging to the Ras superfamily, which has been implicated in AD.^48^ RhoA regulates cytoskeletal rearrangement controlling various neuronal functions. In AD, RhoA levels are diminished overall in the hippocampus.^49^ Rho may mediate neurite retraction^49^ and dendritic spine loss,^50^ possibly by interacting with Aβ^51^ and tau,^52^ overall suggesting putative RhoA participation in neurodegeneration. Similarly, MAPK signaling is a network that amplifies molecules promoting cell proliferation, growth, and survival.^53^ Several MAPK signaling member levels are altered in AD, *e.g*., p38,^54^ JNK,^55^ and their blockade alters amyloid burden. *Mapk8* (JNK) was a DEG in our early-stage Vglut-positive dataset.

Several recurrent themes emerged when we evaluated neuron-specific changes in AD versus control at late-stage disease. Within Pv-positive interneurons, cellular pathways were present as top hits, “MAPK signaling pathway” and “Ras signaling pathway”; however, in the context of the pathway analysis, they may be construed as cancer signaling pathways since many other processes revolving around cancer were present, such as “renal cell carcinoma”, fifth most significant, and “glioma”, ninth most significant, along with a dozen less significant cancers. In contrast, only a few cancer pathways were present in early-stage Pv-positive interneurons and were not among top hits. The involvement of conventional oncogenic pathways are increasingly investigated in neurodegenerative disease, including AD.^53^ In addition to MAPK and Ras, “PI3K-Akt signaling pathway” and “mTOR signaling pathway”, along with DEGs *Cdk*, *Tgfb*, *Trp53*, and *Wnt*, were in our dataset and are implicated in cancer and, more recently, in AD.^53^ Neuronal and synapse pathways were also represented in late-stage Pv-positive interneurons along with insulin resistance and several additional hormone pathways. Central insulin resistance in AD is well-documented along with glucose hypometabolism,^56^ and metabolic dysfunction, such as type 2 diabetes is a dementia risk factor,^57^ linked to bioenergetics failure. Late-stage Sst-positive interneurons continued to showcase glycan biosynthesis, albeit less significantly compared to other pathways than at early-stage, along with various infection, immune phagocytosis, hormone, and metabolic, especially nucleotide, pathways.

Interestingly, late-stage excitatory neurons were primarily characterized by neurodegenerative pathways, in contrast to their early-stage counterparts. Moreover, the excitatory neuron gene expression profile contrasts with that of Pv-positive interneurons, which more strongly express a neurodegenerative profile in early- versus late-stage disease. As posited above, it is conceivable this represents temporal involvement in AD pathology, starting in Pv-positive interneurons and spreading to excitatory neurons later in the disease course. In addition, more cancer pathways were also represented in late-stage excitatory neurons versus their early-stage counterparts, as was the case in interneurons. Moreover, several cell signaling and metabolic pathways with homeostatic as well as oncogenic roles were seen in late-stage excitatory neurons, such as “mTOR signaling pathway”, “choline metabolism in cancer”, and “central carbon metabolism in cancer”. Although metabolic pathways were seen in Pv-positive interneurons, “oxidative phosphorylation” was third most significant in late-stage Vglut-positive neurons, with rich factor of 0.6, along with enrichment of “citrate cycle (TCA cycle)”, both centered on mitochondrial metabolism, which is impaired and a hallmark of AD.^56, 58^ These were accompanied by insulin signaling and insulin resistance, as in late-stage Pv-positive interneurons, not present at early-stage.

Finally, DEGs analysis and pathway enrichment was performed in 5XFAD versus control at early- and late-stage disease examined brain region-dependent differences. We were more limited in our ability to detect DEGs and enriched pathways when we further compared candidates by area as well as by stage (early- versus late-) and neuron type. Many pathways among the most significant were involved in neurodegeneration, cancer, and metabolism. Generally, within each neuron subtype, zero or few DEGs overlapped across all regions, and proportionately few compared to total DEGs were between 2 or 3 regions, *e.g*., DG with CA1, or DG with CA1 with CA3. This possibly suggests interneurons and excitatory neurons mostly displayed distinct profile by brain area.

Our study has several strengths. We examined early- and late-stage disease to identify both initiating or contributing events as well as markers of progressive illness. Moreover, selecting a spatial technology allowed us to profile gene expression *in situ* and isolate ROIs of Pv- and Sst-immunoreactivity to examine interneuron-specific transcriptomics changes, despite that they are far outnumbered by excitatory neurons.^29^ Our study also had weaknesses. We only used male mice limiting our ability to generalize our findings. Additionally, by including excitatory neurons in our analysis, as a counterpoint for comparison, we had a lower resolution for detecting DEGs, and consequently pathways, in interneurons since they constitute a smaller fraction of brain neurons.^29^ Starting with this global analysis of all DEGs in all neurons types led to multiple comparisons across three times more DEGs, restricting significance by adjusted p-value.

In summary, global comparison showed distinct biological pathways related to RNA and protein processing, transport, and clearance in early-stage disease in AD versus control with neurodegeneration pathways at late-stage disease. Examination by neuron subtype suggests possible neurodegenerative pathways emerge within Pv-positive interneurons before excitatory neurons and may possibly reflect a temporal trend in neurodegeneration progression. Moreover, oncogenic pathways with implications in neurodegenerative disease tended to emerge in all neuron subtypes in late-stage. Overall, herein, we present fine-grained transcriptomic profiles for Pv- and Sst-positive interneurons versus Vglut-positive excitatory neurons in a time- and spatial-dependent manner, offering insight into potential AD pathophysiology and therapeutic targets.

## ACKNOWLEDGMENTS

The authors acknowledge the University of Michigan Advanced Genomics Core. This research was supported by the National Institutes of Health (U01AG057562 to ELF; K99AG081390 to MHN), the Alzheimer’s Association (AACSF-22-970586 to KSC), The Robert E. Nederlander Sr. Program for Alzheimer’s Research (ELF), The Andrea and Lawrence A. Wolfe Brain Health Initiative (ELF), The Frank L. and Helen Gofrank Foundation Research Program in Alzheimer’s Disease and Brain Health (ELF), The Sinai Medical Staff Foundation (ELF), The Handleman Emerging Scholar Program (KSC), and The NeuroNetwork for Emerging Therapies (ELF).

## AUTHOR CONTRIBUTIONS

Conceptualization KSC, ELF; Methodology KSC, MHN, ELF; Software MHN; Formal analysis MHN; Investigation KSC, DMR, JMH; Writing - Original Draft MGS, ELF; Writing - Review & Editing all authors; Visualization MHN, MGS; Supervision KSC, ELF; Project administration ELF; Funding acquisition KSC, ELF.

## SUPPLEMENTARY TABLE LEGENDS

**Supplementary Table S1. Global differentially expressed genes (DEGs) in AD (5XFAD) versus control brain at early- and late-stage disease**. DEGs (gene) by brain region and neuron type [parvalbumin (pvalb)-positive interneuron, somatostatin (sst)-positive interneuron, vesicular glutamate transporter (vglut)-positive excitatory neuron] that are shared between early- and late-stage (sheet 1), unique to early-stage (sheet 2), and unique to late-stage (sheet 3) disease. Padj, adjusted p-value; Ctx, cortex; DG, dentate gyrus.

**Supplementary Table S2. KEGG pathway enrichment of AD (5XFAD) versus control brain at early- and late-stage disease**. Kyoto Encyclopedia of Genes and Genomes (KEGG) pathway enrichment of differentially expressed genes (DEGs) shared between early- and late-stage (sheet 1), unique to early-stage (sheet 2), and unique to late-stage (sheet 3) disease in AD (5XFAD) versus control brain. Annot, pathway ID from the database; Term, KEGG pathway name; Annotated, number of DEGs within KEGG pathway; Significant, number of significant DEGs within KEGG pathway; P-value, significance level KEGG pathway; Padj, adjusted significance level KEGG pathway; GeneID, gene name for KEGG pathway DEGs.

**Supplementary Table S3. GO pathway enrichment of AD (5XFAD) versus control brain at early- and late-stage disease**. Gene Ontology (GO) pathway enrichment of differentially expressed genes (DEGs) shared between early- and late-stage (sheet 1), unique to early-stage (sheet 2), and unique to late-stage (sheet 3) disease in AD (5XFAD) versus control brain. Annot, pathway ID from the database; Term, GO pathway name; Annotated, number of DEGs within GO pathway; Significant, number of significant DEGs within GO pathway; P-value, significance level GO pathway; Padj, adjusted significance level GO pathway; GeneID, gene name for GO pathway DEGs.

**Supplementary Table S4. Interneuron-specific differentially expressed genes (DEGs) in AD (5XFAD) versus control brain at early-stage disease**. DEGs (gene) in parvalbumin (Pv)- positive interneurons (sheet 1), somatostatin (Sst)-positive interneurons (sheet 2), and vesicular glutamate transporter (Vglut)-positive excitatory neurons (sheet 3) in AD (5XFAD) versus control at early-stage. Sheets 1 and 2 ordered by lowest to highest p-value; sheet 3 ordered by lowest to highest adjusted p-value (Padj). Ctx, cortex.

**Supplementary Table S5. Interneuron-specific KEGG pathway enrichment of AD (5XFAD) versus control brain at early-stage disease**. Kyoto Encyclopedia of Genes and Genomes (KEGG) pathway enrichment of differentially expressed genes (DEGs) unique to parvalbumin (Pv)-positive interneurons (sheet 1), somatostatin (Sst)-positive interneurons (sheet 2), and vesicular glutamate transporter (Vglut)-positive excitatory neurons (sheet 3) in AD (5XFAD) versus control at early-stage. Annot, pathway ID from the database; Term, KEGG pathway name; Annotated, number of DEGs within KEGG pathway; Significant, number of significant DEGs within KEGG pathway; P-value, significance level KEGG pathway; Padj, adjusted significance level KEGG pathway; GeneID, gene name for KEGG pathway DEGs.

**Supplementary Table S6. Interneuron-specific GO pathway enrichment of AD (5XFAD) versus control brain at early-stage disease**. Gene Ontology (GO) pathway enrichment of differentially expressed genes (DEGs) unique to parvalbumin (Pv)-positive interneurons (sheet 1), somatostatin (Sst)-positive interneurons (sheet 2), and vesicular glutamate transporter (Vglut)-positive excitatory neurons (sheet 3) in AD (5XFAD) versus control at early-stage. Annot, pathway ID from the database; Term, GO pathway name; Annotated, number of DEGs within GO pathway; Significant, number of significant DEGs within GO pathway; P-value, significance level GO pathway; Padj, adjusted significance level GO pathway; GeneID, gene name for GO pathway DEGs.

**Supplementary Table S7. Interneuron-specific differentially expressed genes (DEGs) in AD (5XFAD) versus control brain at late-stage disease**. DEGs (gene) in parvalbumin (Pv)-positive interneurons (sheet 1), somatostatin (Sst)-positive interneurons (sheet 2), and vesicular glutamate transporter (Vglut)-positive excitatory neurons (sheet 3) in AD (5XFAD) versus control at late-stage. All sheets ordered by lowest to highest adjusted p-value (Padj). Ctx, cortex.

**Supplementary Table S8. Interneuron-specific KEGG pathway enrichment of AD (5XFAD) versus control brain at late-stage disease**. Kyoto Encyclopedia of Genes and Genomes (KEGG) pathway enrichment of differentially expressed genes (DEGs) unique to parvalbumin (Pv)-positive interneurons (sheet 1), somatostatin (Sst)-positive interneurons (sheet 2), and vesicular glutamate transporter (Vglut)-positive excitatory neurons (sheet 3) in AD (5XFAD) versus control at late-stage. Annot, pathway ID from the database; Term, KEGG pathway name; Annotated, number of DEGs within KEGG pathway; Significant, number of significant DEGs within KEGG pathway; P-value, significance level KEGG pathway; Padj, adjusted significance level KEGG pathway; GeneID, gene name for KEGG pathway DEGs.

**Supplementary Table S9. Interneuron-specific GO pathway enrichment of AD (5XFAD) versus control brain at late-stage disease**. Gene Ontology (GO) pathway enrichment of differentially expressed genes (DEGs) unique to parvalbumin (Pv)-positive interneurons (sheet 1), somatostatin (Sst)-positive interneurons (sheet 2), and vesicular glutamate transporter (Vglut)-positive excitatory neurons (sheet 3) in AD (5XFAD) versus control at late-stage. Annot, pathway ID from the database; Term, GO pathway name; Annotated, number of DEGs within GO pathway; Significant, number of significant DEGs within GO pathway; P-value, significance level GO pathway; Padj, adjusted significance level GO pathway; GeneID, gene name for GO pathway DEGs.

**Supplementary Table S10. Pv-positive interneuron differentially expressed genes (DEGs) in AD (5XFAD) versus control brain regions at early-stage disease**. DEGs (gene) in parvalbumin (Pv)-positive interneurons in dentate gyrus (DG, sheet 1), CA1 (sheet 2), CA3 (sheet 3), and cortex (ctx, sheet 4) in AD (5XFAD) versus control at early-stage. All sheets ordered by lowest to highest p-value. Padj, adjusted p-value.

**Supplementary Table S11. Pv-positive interneuron KEGG pathway enrichment of AD (5XFAD) versus control brain regions at early-stage disease**. Kyoto Encyclopedia of Genes and Genomes (KEGG) pathway enrichment of differentially expressed genes (DEGs) unique to parvalbumin (Pv)-positive interneurons in dentate gyrus (DG, sheet 1), CA1 (sheet 2), CA3 (sheet 3), and cortex (sheet 4) in AD (5XFAD) versus control at early-stage. Annot, pathway ID from the database; Term, KEGG pathway name; Annotated, number of DEGs within KEGG pathway; Significant, number of significant DEGs within KEGG pathway; P-value, significance level KEGG pathway; Padj, adjusted significance level KEGG pathway; GeneID, gene name for KEGG pathway DEGs.

**Supplementary Table S12. Pv-positive interneuron GO pathway enrichment of AD (5XFAD) versus control brain regions at early-stage disease**. Gene Ontology (GO) pathway enrichment of differentially expressed genes (DEGs) unique to parvalbumin (Pv)-positive interneurons in dentate gyrus (DG, sheet 1), CA1 (sheet 2), CA3 (sheet 3), and cortex (sheet 4) in AD (5XFAD) versus control at early-stage. Annot, pathway ID from the database; Term, GO pathway name; Annotated, number of DEGs within GO pathway; Significant, number of significant DEGs within GO pathway; P-value, significance level GO pathway; Padj, adjusted significance level GO pathway; GeneID, gene name for GO pathway DEGs.

**Supplementary Table S13. Sst-positive interneuron differentially expressed genes (DEGs) in AD (5XFAD) versus control brain regions at early-stage disease**. DEGs (gene) in somatostatin (Sst)-positive interneurons in dentate gyrus (DG, sheet 1), CA1 (sheet 2), CA3 (sheet 3), and cortex (ctx, sheet 4) in AD (5XFAD) versus control at early-stage. All sheets ordered by lowest to highest p-value. Padj, adjusted p-value.

**Supplementary Table S14. Sst-positive interneuron KEGG pathway enrichment of AD (5XFAD) versus control brain regions at early-stage disease**. Kyoto Encyclopedia of Genes and Genomes (KEGG) pathway enrichment of differentially expressed genes (DEGs) unique to somatostatin (Sst)-positive interneurons in dentate gyrus (DG, sheet 1), CA1 (sheet 2), CA3 (sheet 3), and cortex (sheet 4) in AD (5XFAD) versus control at early-stage. Annot, pathway ID from the database; Term, KEGG pathway name; Annotated, number of DEGs within KEGG pathway; Significant, number of significant DEGs within KEGG pathway; P-value, significance level KEGG pathway; Padj, adjusted significance level KEGG pathway; GeneID, gene name for KEGG pathway DEGs.

**Supplementary Table S15. Sst-positive interneuron GO pathway enrichment of AD (5XFAD) versus control brain regions at early-stage disease**. Gene Ontology (GO) pathway enrichment of differentially expressed genes (DEGs) unique to somatostatin (Sst)-positive interneurons in dentate gyrus (DG, sheet 1), CA1 (sheet 2), CA3 (sheet 3), and cortex (sheet 4) in AD (5XFAD) versus control at early-stage. Annot, pathway ID from the database; Term, GO pathway name; Annotated, number of DEGs within GO pathway; Significant, number of significant DEGs within GO pathway; P-value, significance level GO pathway; Padj, adjusted significance level GO pathway; GeneID, gene name for GO pathway DEGs.

**Supplementary Table S16. Vglut-positive interneuron differentially expressed genes (DEGs) in AD (5XFAD) versus control brain regions at early-stage disease**. DEGs (gene) in vesicular glutamate transporter (Vglut)-positive interneurons in dentate gyrus (DG, sheet 1), CA1 (sheet 2), CA3 (sheet 3), and cortex (ctx, sheet 4) in AD (5XFAD) versus control at early-stage. Sheets 3 and 4 ordered by lowest to highest p-value; sheets 1 and 2 ordered by lowest to highest adjusted p-value (Padj).

**Supplementary Table S17. Vglut-positive interneuron KEGG pathway enrichment of AD (5XFAD) versus control brain regions at early-stage disease**. Kyoto Encyclopedia of Genes and Genomes (KEGG) pathway enrichment of differentially expressed genes (DEGs) unique to vesicular glutamate transporter (Vglut)-positive interneurons in dentate gyrus (DG, sheet 1), CA1 (sheet 2), CA3 (sheet 3), and cortex (sheet 4) in AD (5XFAD) versus control at early-stage. Annot, pathway ID from the database; Term, KEGG pathway name; Annotated, number of DEGs within KEGG pathway; Significant, number of significant DEGs within KEGG pathway; P-value, significance level KEGG pathway; Padj, adjusted significance level KEGG pathway; GeneID, gene name for KEGG pathway DEGs.

**Supplementary Table S18. Vglut-positive interneuron GO pathway enrichment of AD (5XFAD) versus control brain regions at early-stage disease**. Gene Ontology (GO) pathway enrichment of differentially expressed genes (DEGs) unique to vesicular glutamate transporter (Vglut)-positive interneurons in dentate gyrus (DG, sheet 1), CA1 (sheet 2), CA3 (sheet 3), and cortex (sheet 4) in AD (5XFAD) versus control at early-stage. Annot, pathway ID from the database; Term, GO pathway name; Annotated, number of DEGs within GO pathway; Significant, number of significant DEGs within GO pathway; P-value, significance level GO pathway; Padj, adjusted significance level GO pathway; GeneID, gene name for GO pathway DEGs.

**Supplementary Table S19. Pv-positive interneuron differentially expressed genes (DEGs) in AD (5XFAD) versus control brain regions at late-stage disease**. DEGs (gene) in parvalbumin (Pv)-positive interneurons in dentate gyrus (DG, sheet 1), CA1 (sheet 2), CA3 (sheet 3), and cortex (ctx, sheet 4) in AD (5XFAD) versus control at late-stage. Sheets 1 and 2 ordered by lowest to highest p-value; sheets 3 and 4 ordered by lowest to highest adjusted p-value (Padj).

**Supplementary Table S20. Pv-positive interneuron KEGG pathway enrichment of AD (5XFAD) versus control brain regions at late-stage disease**. Kyoto Encyclopedia of Genes and Genomes (KEGG) pathway enrichment of differentially expressed genes (DEGs) unique to parvalbumin (Pv)-positive interneurons in dentate gyrus (DG, sheet 1), CA1 (sheet 2), CA3 (sheet 3), and cortex (sheet 4) in AD (5XFAD) versus control at late-stage. Annot, pathway ID from the database; Term, KEGG pathway name; Annotated, number of DEGs within KEGG pathway; Significant, number of significant DEGs within KEGG pathway; P-value, significance level KEGG pathway; Padj, adjusted significance level KEGG pathway; GeneID, gene name for KEGG pathway DEGs.

**Supplementary Table S21. Pv-positive interneuron GO pathway enrichment of AD (5XFAD) versus control brain regions at late-stage disease**. Gene Ontology (GO) pathway enrichment of differentially expressed genes (DEGs) unique to parvalbumin (Pv)-positive interneurons in dentate gyrus (DG, sheet 1), CA1 (sheet 2), CA3 (sheet 3), and cortex (sheet 4) in AD (5XFAD) versus control at late-stage. Annot, pathway ID from the database; Term, GO pathway name; Annotated, number of DEGs within GO pathway; Significant, number of significant DEGs within GO pathway; P-value, significance level GO pathway; Padj, adjusted significance level GO pathway; GeneID, gene name for GO pathway DEGs.

**Supplementary Table S22. Sst-positive interneuron differentially expressed genes (DEGs) in AD (5XFAD) versus control brain regions at late-stage disease**. DEGs (gene) in somatostatin (Sst)-positive interneurons in dentate gyrus (DG, sheet 1), CA1 (sheet 2), CA3 (sheet 3), and cortex (ctx, sheet 4) in AD (5XFAD) versus control at late-stage. Sheets 1, 2, and 4 ordered by lowest to highest p-value; sheet 3 ordered by lowest to highest adjusted p-value (Padj).

**Supplementary Table S23. Sst-positive interneuron KEGG pathway enrichment of AD (5XFAD) versus control brain regions at late-stage disease**. Kyoto Encyclopedia of Genes and Genomes (KEGG) pathway enrichment of differentially expressed genes (DEGs) unique to somatostatin (Sst)-positive interneurons in dentate gyrus (DG, sheet 1), CA1 (sheet 2), CA3 (sheet 3), and cortex (sheet 4) in AD (5XFAD) versus control at late-stage. Annot, pathway ID from the database; Term, KEGG pathway name; Annotated, number of DEGs within KEGG pathway; Significant, number of significant DEGs within KEGG pathway; P-value, significance level KEGG pathway; Padj, adjusted significance level KEGG pathway; GeneID, gene name for KEGG pathway DEGs.

**Supplementary Table S24. Sst-positive interneuron GO pathway enrichment of AD (5XFAD) versus control brain regions at late-stage disease. Gene Ontology (GO) pathway**

enrichment of differentially expressed genes (DEGs) unique to somatostatin (Sst)-positive interneurons in dentate gyrus (DG, sheet 1), CA1 (sheet 2), CA3 (sheet 3), and cortex (sheet 4) in AD (5XFAD) versus control at late-stage. Annot, pathway ID from the database; Term, GO pathway name; Annotated, number of DEGs within GO pathway; Significant, number of significant DEGs within GO pathway; P-value, significance level GO pathway; Padj, adjusted significance level GO pathway; GeneID, gene name for GO pathway DEGs.

**Supplementary Table S25. Vglut-positive interneuron differentially expressed genes (DEGs) in AD (5XFAD) versus control brain regions at late-stage disease**. DEGs (gene) in vesicular glutamate transporter (Vglut)-positive interneurons in dentate gyrus (DG, sheet 1), CA1 (sheet 2), CA3 (sheet 3), and cortex (ctx, sheet 4) in AD (5XFAD) versus control at late-stage. All sheets ordered by lowest to highest adjusted p-value (Padj).

**Supplementary Table S26. Vglut-positive interneuron KEGG pathway enrichment of AD (5XFAD) versus control brain regions at late-stage disease**. Kyoto Encyclopedia of Genes and Genomes (KEGG) pathway enrichment of differentially expressed genes (DEGs) unique to vesicular glutamate transporter (Vglut)-positive interneurons in dentate gyrus (DG, sheet 1), CA1 (sheet 2), CA3 (sheet 3), and cortex (sheet 4) in AD (5XFAD) versus control at late-stage. Annot, pathway ID from the database; Term, KEGG pathway name; Annotated, number of DEGs within KEGG pathway; Significant, number of significant DEGs within KEGG pathway; P-value, significance level KEGG pathway; Padj, adjusted significance level KEGG pathway; GeneID, gene name for KEGG pathway DEGs.

**Supplementary Table S27. Vglut-positive interneuron GO pathway enrichment of AD (5XFAD) versus control brain regions at late-stage disease**. Gene Ontology (GO) pathway enrichment of differentially expressed genes (DEGs) unique to vesicular glutamate transporter (Vglut)-positive interneurons in dentate gyrus (DG, sheet 1), CA1 (sheet 2), CA3 (sheet 3), and cortex (sheet 4) in AD (5XFAD) versus control at late-stage. Annot, pathway ID from the database; Term, GO pathway name; Annotated, number of DEGs within GO pathway; Significant, number of significant DEGs within GO pathway; P-value, significance level GO pathway; Padj, adjusted significance level GO pathway; GeneID, gene name for GO pathway DEGs.

---

**Supplementary Figure S1.**
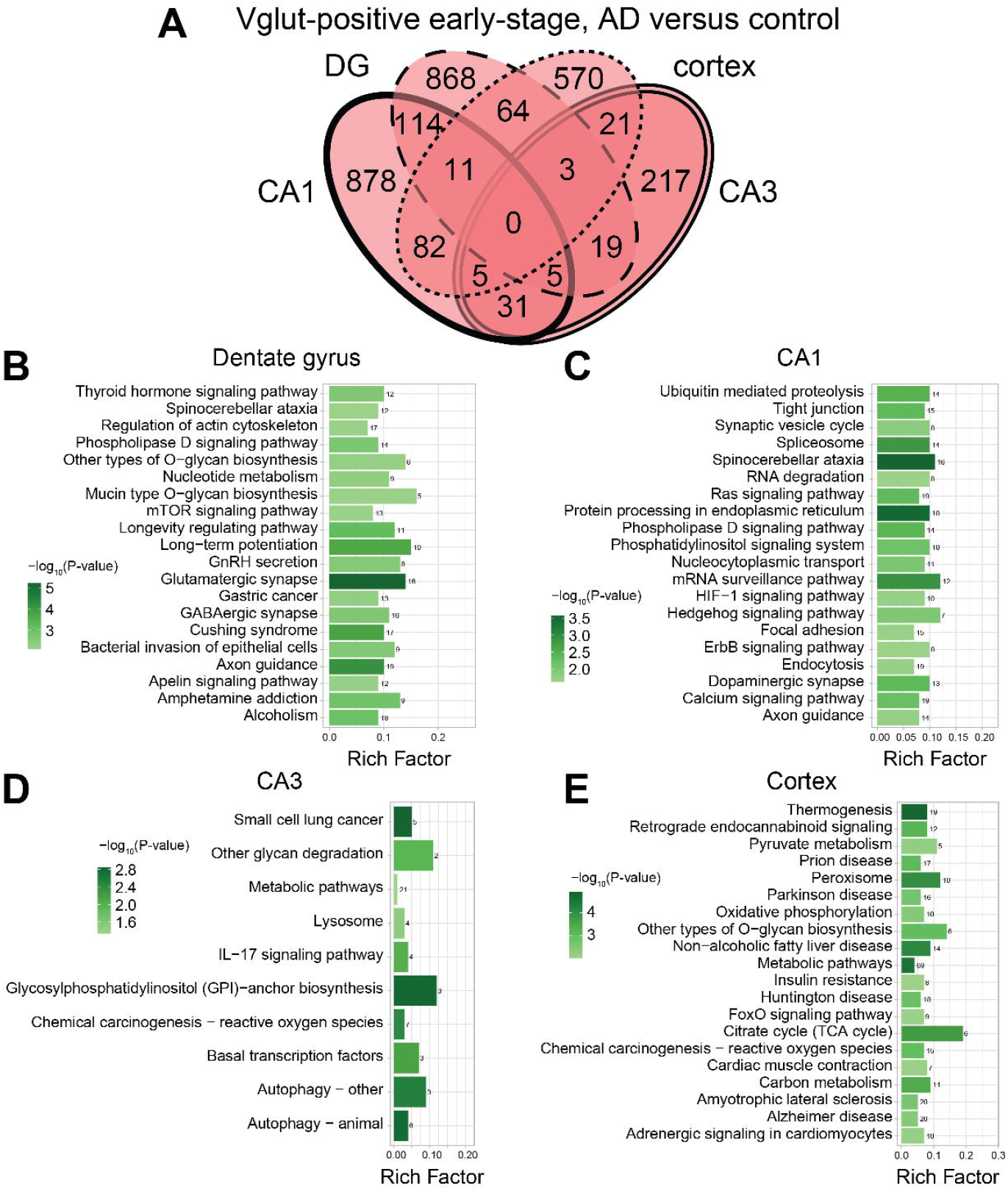
Region-specific DEGs in AD (5XFAD) versus control Vglut- positive excitatory neurons at early-stage disease. (**A**) Venn diagram of differentially expressed genes (DEGs) in AD (5XFAD) versus control across all analyzed regions dentate gyrus (DG), CA1, CA3, and cortex in Vglut-positive excitatory neurons at early-stage disease. KEGG pathway enrichment analysis of early-stage DEGs in 5XFAD AD versus control Vglut- positive excitatory neurons in (**B**) DG (dashed), (**C**) CA1 (bold), (**D**) CA3 (dotted), (**E**) cortex (double-line). Legend represents significance level by -log_10_(p-value), numbers over the bars represent number of DEGs belonging to annotated pathways. Background non-neuronal DEGs were subtracted from the analysis.

**Supplementary Figure S2.**
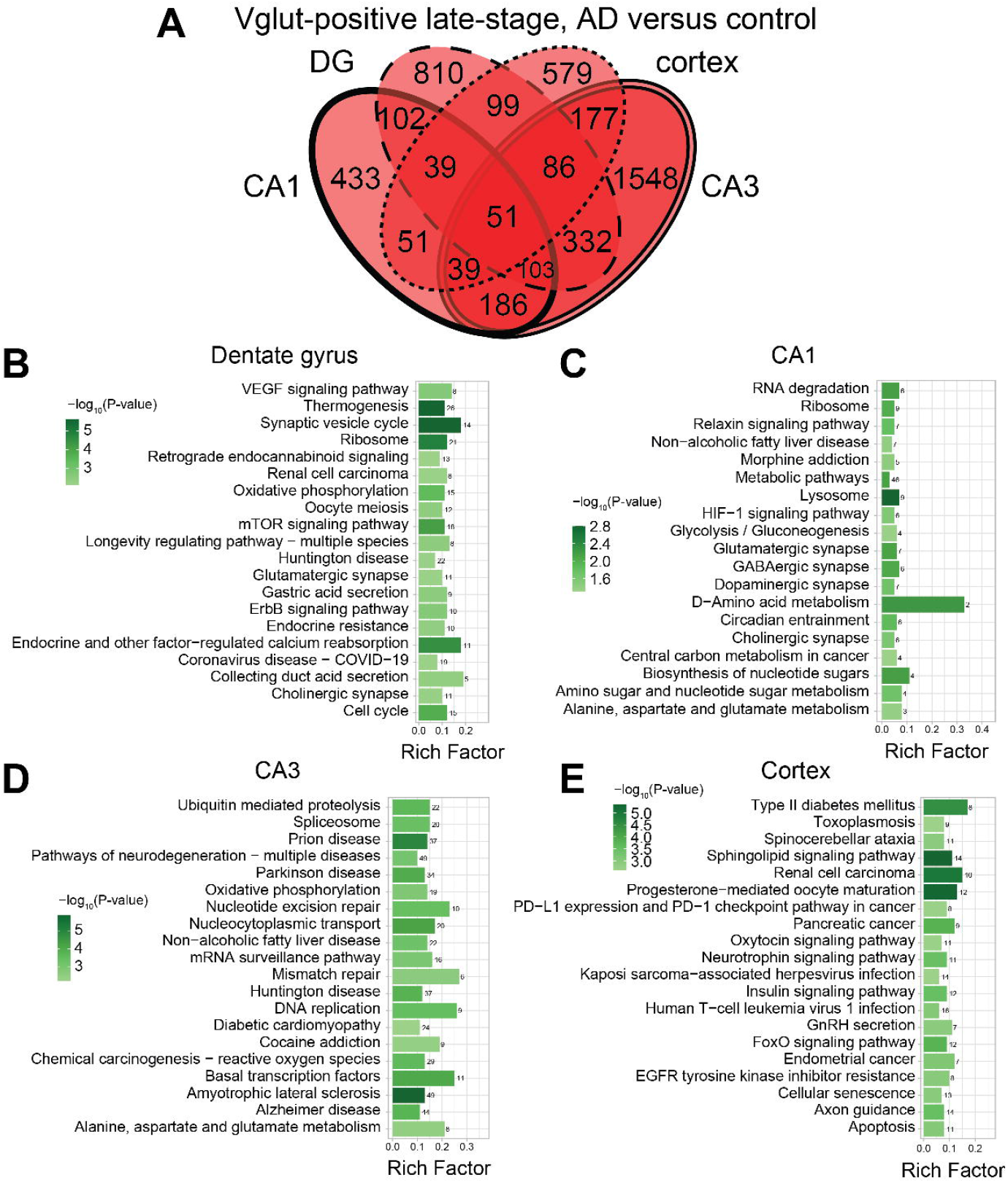
Region-specific DEGs in AD (5XFAD) versus control Vglut- positive excitatory neurons at late-stage disease. (**A**) Venn diagram of differentially expressed genes (DEGs) in AD (5XFAD) versus control across all analyzed regions dentate gyrus (DG), CA1, CA3, and cortex in Vglut-positive excitatory neurons at late-stage disease. KEGG pathway enrichment analysis of late-stage DEGs in 5XFAD AD versus control Vglut- positive excitatory neurons in (**B**) DG (dashed), (**C**) CA1 (bold), (**D**) CA3 (dotted), (**E**) cortex (double-line). Legend represents significance level by -log_10_(p-value), numbers over the bars represent number of DEGs belonging to annotated pathways. Background non-neuronal DEGs were subtracted from the analysis.

